# EBV Triggers a Distinct Antiviral Response in HMC3 Cells

**DOI:** 10.64898/2026.04.03.716358

**Authors:** Noah E. Berkowitz, Alexander Nosov, Mark Nosov, Fryda Solis Roldan, Anuj Ahuja, Miranda McGaskey, Ethel Cesarman, Douglas F. Nixon, Nicholas Dopkins

**Affiliations:** Northwell Health, New Hyde Park, NY, USA; Feinstein Institutes for Medical Research, Manhasset, NY, USA; Department of Pathology and Laboratory Medicine, Weill Cornell Medicine, New York, New York, USA; Zucker School of Medicine at Hofstra/Northwell-Hofstra University, Hempstead, NY, USA

## Abstract

Epstein-Barr Virus (EBV) is a gamma herpesvirus found in >90% of the world population that is associated with primary central nervous system (CNS) malignancy development in immunocompromised people. To provide mechanistic links between EBV infection and CNS malignancies, we investigated the capacity for EBV particles to suppress anti-tumor immunity in human microglia through a cell line model. With this approach, we exposed HMC3 cells to EBV-derived glycoprotein 350 (GP350), UV-inactivated EBV (UVi-EBV), and lipoteichoic acid (LTA) for up to 72 hours. Acute impacts of EBV particles and glycoprotein on microglial physiology were characterized at various timepoints in this model through measures of cytokine production, mRNA expression, and endocytosis. We found that UVi-EBV exposure significantly suppressed microglial production of anti-tumor interferons (IFNs) and upregulated microglial expression of the proto-oncogenic immediate early genes FOS and EGR1. Notably, there was no impairment of microglial endocytic functions following UVi-EBV stimulation, suggesting a compartmentalized suppression on IFN signaling. Overall, these findings reveal that the EBV-mediated inhibition of microglial IFN production may contribute to CNS malignancies and emphasize the urgency of innovating therapeutic strategies which target EBV to restore microglial anti-tumoral immunity.

**Importance:** Evidence linking EBV infection with primary CNS lymphomas and leiomyosarcomas are abundant, yet it is unclear whether EBV infection influences the CNS microenvironment and whether these effects then promote tumorigenesis. This study demonstrates evidence for EBV particle exposure to influence microglial immune phenotypes by suppressing IFN production, providing a putative mechanism for EBV virion expression in the CNS to suppress anti-tumoral immunity against EBV+ cancers. These results are particularly relevant to the etiology of EBV+ primary CNS cancers in immunocompromised people, where microglial play a heightened role in protecting the CNS in the absence of adaptive immunity.

## 1. Introduction

Epstein-Barr Virus (EBV) is highly infectious gamma herpesvirus found in >90% of the population(1). EBV is transmission occurs via oropharyngeal secretions as the virus infect the oral mucosa epithelium before transmitting to proximal naïve B cells beneath or within the mucosal barrier(2, 3). Infected memory B cells then harbor long lasting reservoirs of latent EBV(4). B cell reservoirs may be driven to resume lytic replication and virion production by external factors such as stress or immune suppression(4, 5). Sustained mononucleosis and latent EBV reservoirs frequently disrupt host immunity and B cell biology(6), putatively linking EBV infections with the later onset and severity of chronic diseases, such as inflammatory disorders and lymphoproliferative malignancies(7–9). Overwhelming associations between EBV infection status and cancers (e.g., lymphomas, carcinomas, and sarcomas) have designated the virus as a tumor promoting class 1 carcinogen(1, 10, 11). Despite exhaustive efforts demonstrating how EBV directly, or indirectly, shapes tumorigenic landscapes(12, 13), it remains unclear why, and how, EBV infection is carcinogenic in some, but not all, of the infected population. Cofactors stemming from genetics, immunocompetency, age, co-infections, gender, environment, viral latency stages, and background may likely confer resistances or susceptibilities to EBV-driven effects on the body and therefore require thorough investigation(9, 14–20).

Immunodeficiencies render individuals less capable, or entirely incapable, of fighting infections, such as EBV(21). Immunodeficiencies may result from genetic factors (primary immunodeficiencies), infections such as with HIV, hematological malignancies, or from pharmacological or radiological depletion of the immune system to combat autoimmune inflammation or prepare for transplantation procedures (secondary immunodeficiencies)(22). Retrospective estimates suggest that ∼1.8-1.9% of Canadian, ∼6.6% of U.S, and ∼2.6% of Chinese populations exhibit varying levels of immunodeficiency and that their status rates are rising(22–24). EBV infection in immunocompromised individuals can be especially detrimental to the immunologically unique and privileged microenvironments which comprise the central nervous system (CNS). Aggressive and rare cancers such as primary CNS lymphomas (PCNSLs) and primary intracranial leiomyosarcomas (PILMs) typically only form as EBV+ tumors in severely immunocompromised individuals(10, 25–29). EBV infection has also been implicated as a major risk factor for multiple sclerosis(30), an autoimmune disorder of the CNS in which the immune system recognizes peptides within the myelin sheath protecting axons(31, 32). However, the pleiotropic mechanisms by which EBV infection may simultaneously contribute to an immunosuppressive tumor microenvironment (TME) in PCNSLs/PILMs and proinflammatory plaque formation in MS remain unclear and requires further study.

As the incidence rates for inflammatory disorders and compromised immunity rise, we argue that characterizing the impact of EBV infection on the neuroimmune niche(s) of the CNS is of importance. For this purpose, we investigated whether EBV particle exposure influences immune phenotypes in a cell line model of human microglia, the CNS-resident mononuclear phagocytes that play key roles in maintaining CNS plasticity and sterility(33). Microglia effectively prime anti-tumor immunity in the CNS by recognizing cancer-associated molecular patterns to stimulate interferon (IFN) production and activate tumor reactive lymphocytes(34, 35).

The role for microglial cells to protect the CNS from cancerous cells would also be heightened in the absence of competent adaptive immune responses, providing putative roles for EBV-microglial interactions to facilitate PCNSL/PILM generation in severely immunocompromised people(36). Based on our understanding of neuroimmune dynamics in immunocompromised persons and associations with EBV+ primary CNS tumor incidence, we hypothesized that EBV promotes an immunoregulatory bystander phenotype in microglial cells that is conducive to primary tumorigenesis(37). To evaluate this hypothesis, we utilized the HMC3 cell line as an *in vitro* model of human microglia to stimulate with either EBV viral glycoprotein (GP350), UV-inactivated EBV (UVi-EBV), or lipoteichoic acid (LTA). LTA, a TLR2 agonist, was used as a positive control and possible route for examining TLR2-mediated recognition of EBV dUTPase(38). Microglia phenotypes were then characterized over a time period of 72-hours by measuring cytokine production levels and endocytosis with the underlying transcriptional changes analyzed via RNA sequencing. Ultimately, this work aims to elucidate how a viral pathogen disrupts the neuroimmune landscape to participate in CNS disease etiologies.

## 2. Methods

### 2.1 HMC3 Cell Culturing

Purified EBV virions were obtained from the supernatant of Akata cells cultured within RPMI medium (Hyclone, Cat#SH30027.01) containing 10% fetal bovine serum (FBS; Gibco, Cat#A52567-01) and 1% penicillin/streptomycin (pen/strep; Gibco, Cat#15070063), hereafter referred to as complete RPMI (cRPMI). EBV virion production by Akata cells was induced by treatment with 50 µg/mL anti-human immunoglobulin G (MP Biomedicals, Cat#08634811). Encapsulated EBV genome copies were quantified from DNase treated Akata cell culture supernatants using real-time PCR. Primers (forward: 5′-CCCAACACTCCACCACACC-3′; reverse: 5′-TCTTAGGAGCTGTCCGAGGG-3′) and a probe (5′-[6-FAM]CACACACTACACACACCCACCCGTCTC[BHQ-1]-3′) specific for the BamHI-W region of the EBV genome were used for real-time PCR quantification. Stocks of cRPMI medium were determined to contain 6.4 × 10^9^ genomic copies/mL. Following quantification, EBV particles were inactivated using a Stratalinker 1800 UV crosslinker using an energy dose of 2,500 µJ/cm^2^ × 100 applied five times with 2-minute intervals between exposure.

Human microglial HMC3 cells (ATCC, Cat#CRL-3304) were cultured in EMEM medium (ATCC, Cat#30-2003) supplemented with 10% FBS and 1% pen/strep (Gibco, Cat#15140-122), hereafter referred to as complete EMEM (cEMEM). Cells were maintained in a humidified incubator at 37°C and 5% CO2 and were routinely maintained at 80-90% confluency. All experiments were performed with HMC3 cells from their first passage, which tested negative for mycoplasma contamination. HMC3 cells were seeded into 48-well plates at a density of 10^6^ live cells/mL in cEMEM and allowed to adhere for 72 hours. The spent medium was then replaced with fresh cEMEM containing one of four conditions: Vehicle (VEH; cEMEM w/10% cRPMI v/v; Gibco, Cat#11875-093), GP350 (ACROBiosystems, Cat#GP0-E52H6; 80 ng/mL in cEMEM w/10% cRPMI v/v), LTA (InvivoGen, Cat#tlrl-pslta1; 1μg/mL in cEMEM w/10% cRPMI v/v), and UVi-EBV (6.4 × 10^8^ genomic copies of EBV/mL in cEMEM w/10% cRPMI v/v). At 0-, 2-, 4-, 24-, 48-, and 72-hours post-treatment, culture supernatants were collected and stored at -80°C for cytokine analysis. For RNA collection, adherent cells were washed with PBS (Gibco, Cat#10010-031), harvested by trypsinization, and lysed in Trizol™ reagent (Invitrogen, Cat#15596026). Briefly, cells were detached using 0.05% trypsin-EDTA (Corning, Cat#25-052-CV), the reaction was neutralized with cEMEM, and the cell suspension was pelleted by centrifugation at 500 x g for 5 minutes at 4°C. The resulting cell pellet was lysed in 200 μL of Trizol™ and stored at -80°C.

### 2.2 RNA Extraction and Purification

Total RNA was extracted from Trizol™ lysates using a chloroform (Thermo Scientific, Cat#423555000) phase separation, before being precipitated with isopropanol (Thermo Scientific, Cat#383910010) and being further purified with the columns and washes provided by RNeasy™ Mini Kits (Qiagen, Cat#74104). RNA extracts were then further purified with the RNAClean™ XP Kit (Beckman Coulter, Cat#A63987). RNA integrity was assessed via NanoDrop One™ (Thermo Scientific), RNA Qubit (Thermo Scientific), and TapeStation (Agilent). RNA sequencing was performed at GENEWIZ™ (Southfield, NJ). Poly(A) mRNA was selected using the NEBNext™ Poly(A) Magnetic Isolation Module (New England Biolabs). Libraries were prepared using the NEBNext™ Ultra II RNA Library Prep Kit for Illumina (New England Biolabs), and 150bp paired-end sequencing was performed on an Illumina NovaSeq 6000 instrument™ (Illumina) to a target depth of 30 million reads per sample.

### 2.3 Differential Expression Analysis

FASTQ files were aligned to the human genome build 38 (hg38) using STAR (v2.7.9.a). STAR alignment was performed using the parameters “--outSAMstrandField intronMotif --outFilterMultimapNmax 200 --winAnchorMultimapNmax 200”. Differential expression analysis was performed using standard DESEQ2(39) (v1.30.1) parameters. Results were extracted as DESEQ objects, with a numbered contrast of each group compared against all others. For identifying differentially expressed transcripts, we assumed any tested loci possessing a log2 fold change of >1 or < −1 to indicate biological significance and an adjusted p value (padj) of <0.05 to indicate statistical significance. All monitored loci that surpassing thresholds for biological and statistical significance were then assumed as differentially expressed. Differentially expressed transcripts were visualized with the R package EnhancedVolcano (v1.8.0). Original code for performing downstream analysis in R can be accessed at https://github.com/NicholasDopkins/HMC3.

### 2.4 Cytokine Production Analysis

Cell supernatants were thawed on ice and analyzed using the LEGENDplex™ Human Anti-Virus Response Panel 1 (13-plex) with V-bottom plate (BioLegend, Cat# 740390) according to the manufacturer’s instructions. Samples were analyzed on a NovoCyte Quanteon Flow Cytometer (Agilent) and predicted cytokine concentrations were determined using BioLegend’s LEGENDplex™ Data Analysis Software (BioLegend). Outliers with the predicted concentrations exceeding the highest standard concentration (IL-1β, TNF-α, and IFNβ concentrations for 2-hours GP350 replicate number 1) were removed from downstream analysis due to uncredible cytokine quantifications arising due to human or machine error.

### 2.5 Endocytosis Assay

Endocytosis was measured in in VEH and UVi-EBV treated HMC3 cells using pHrodo™ Green conjugated 10kDa Dextran beads (Invitrogen™, Cat# P10361). Briefly, HMC3 cells were plated at 10^6^ live cells/mL in 12 well plates as previously described. At hour 0, media was exchanged, and cells were exposed to either VEH-or UVi-EBV-exposed conditions. At 0-, 2-, 4-, 24-, 48-, and 72-hours post exposure, the media was decanted, and adherent cells were washed 3 times with PBS. Adherent cells were then incubated with 12.5μg of pHrodo™ Green conjugated 10kDa Dextran beads in 200μL of cEMEM (50μg/mL) for 1 hour. After 1 hour, the media containing pHrodo™ Green conjugated 10kDa Dextran beads were decanted, and adherent cells were washed 3 times with PBS. Cells were then fixed for 30 minutes at room temperature conditions using a fixation buffer containing 4% paraformaldehyde (BioLegend, Cat# 420801). Fixation buffer was then decanted and 200μL of DAPI Fluoromount-G® (SouthernBiotech, Cat# 0100-20) was added to each well for immediate imaging with a Cytation5 instrument using the DAPI and FITC channels. Once imaged, a circle 6,441μm in diameter centered at x=0 and y=0 (center of the well) encompassing a 32.58mm^2^ area was used for analysis. The extent of microglial endocytosis was recorded in the number of endocytic vesicles (EVs) per 1mm^2^ and the sum area of captured EVs per 32.58mm^2^.

### 2.6 Statistics

The default Wald testing parameters within DESEQ2(39) that provide a z score using shrunken estimates of log fold change divided by standard error and a Benjamini-Hochberg correction of p values were utilized for the identification of significantly differential expression of genes between samples. Statistical analysis of cytokine production was performed using a repeated measures ANOVA with the Geisser-Greenhouse correction, followed by Dunnett’s multiple comparison test for the generation of p-values to compare each treatment against the time-matched VEH. For comparisons between samples with unequal grouping due to the removal of cytokine quantification outliers, a mixed effects model with the Geisser-Greenhouse correction followed by Dunnett’s multiple comparison test for the generation of p-values to compare each treatment against the time-matched VEH. Statistical analysis of EV production was performed using a repeated measures two-way ANOVA with the Geisser-Greenhouse correction, followed by Šídák’s multiple comparison test for the generation of p-values to UVi-EBV exposure against the time-matched VEH. Correlations between markers were determined using Spearman’s correlation coefficient. Listed padj are represented to a definition of 5 significant digits. For cytokine profiling, all padj <0.10 are displayed, and padj <0.05 are referred to as significant.

## 3. Results

### UVi-EBV Suppresses Microglial Production of Anti-Tumor Cytokines

To determine the effects of EBV exposure on microglial immune phenotypes, we exposed HMC3 cells to UVi-EBV, GP350, and LTA then quantified cytokine production over 72-hours (Figure 1). Briefly, UVi-EBV exposure significantly inhibited type-I IFNs at 72-hours post treatment, with substantial reductions observed in both IFN-α2 and IFN-β levels (Figure 1B-C). The reduction in type-I IFNs reflect known mechanisms for EBV’s immune evasion, where viral proteins, such as BGLF4 and BPLF1, may block IFN production(40, 41). IFN-γ secretion exhibited a trend of suppression at 72 hours, while IFN-λ2 remained undetectable following UVi-EBV stimulation (Figure 1D-F). The reduction in type-II and type-III IFNs is consistent with results from EBV+ post-transplant lymphoproliferative diseases, where decreased IFN signaling supports immune evasion(3, 42). IP-10, also known as CXCL10, was significantly suppressed at 72-hours following exposure to UVi-EBV particles (Figure 1L), contrasting previous reports of EBV-induced IP-10 expression in B cells and epithelial cells(43), suggesting potential cell-type specific IP-10 responses to EBV. In the TME, the recruitment of cytotoxic T cells can be facilitated by IP-10(44), indicating a mechanism for EBV to impede anti-tumor lymphocyte infiltration in EBV+ PCNSLs and PILMs. Pro-inflammatory cytokines IL-6, IL-8, and TNF-α demonstrated no significant changes with EBV exposure (Figure 1H-I, M), further highlighting the cell-type specific patterns when comparing with the induction of IL-6 and IL-8 expression EBV in nasopharyngeal carcinoma cells(45, 46).

**Figure 1:**
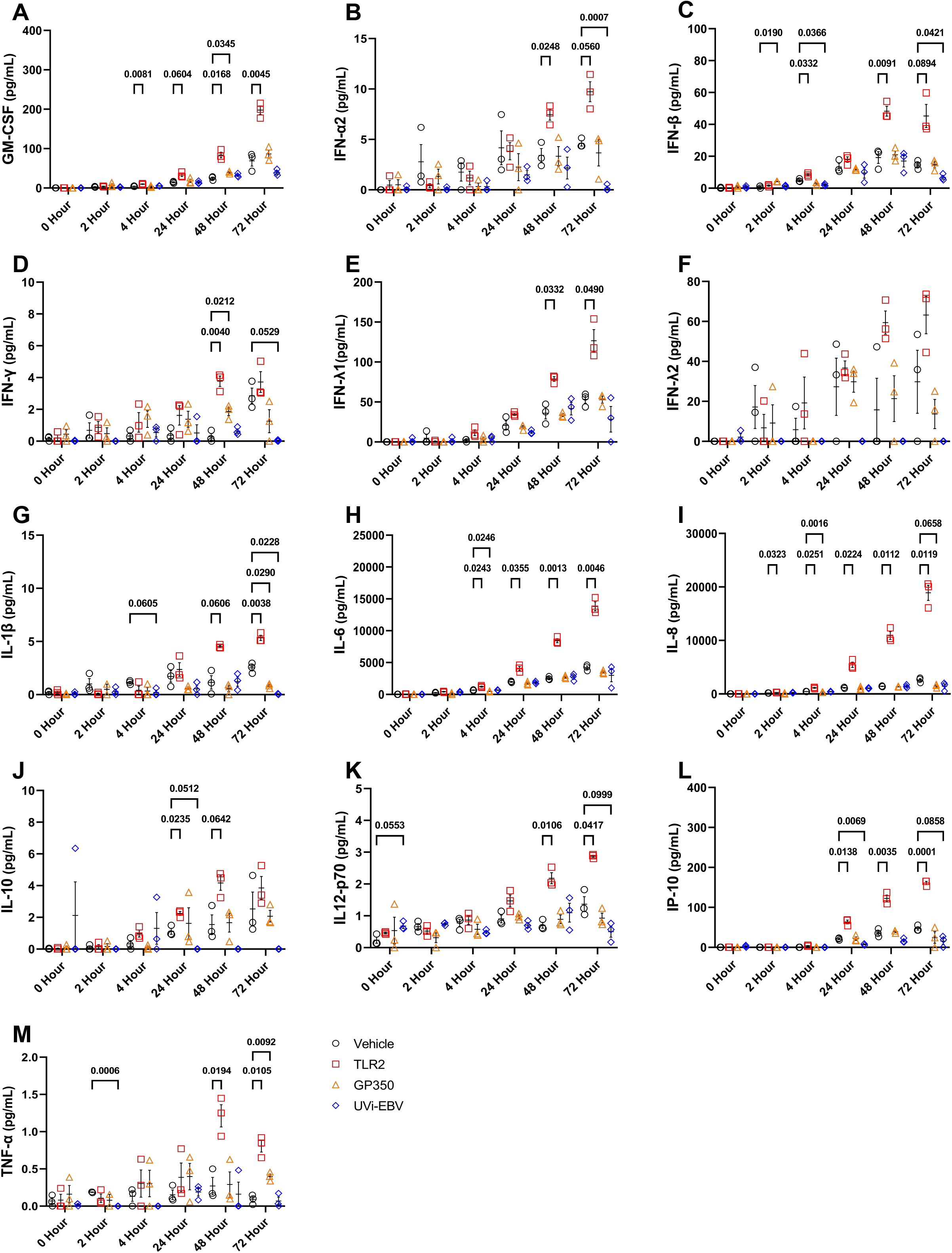
UVi-EBV Suppresses Microglial Production of Anti-Tumor Cytokines. Human microglial HMC3 cells were stimulated with vehicle (VEH; cEMEM w/10% cRPMI), Lipoteichoic Acid (LTA, VEH+1 μg/mL), GP350 (VEH+80 ng/mL), or UV-inactivated EBV (UVi-EBV, VEH+6.4 × 10^8^ genomic copies/mL). Supernatants were collected at 0-, 2-, 4-, 24-, 48-, and 72-hours post-treatment. To control for baseline cytokine levels in the initial UVi-EBV preparation, all data were normalized by subtracting the mean concentration at the 0-hour time point from each ensuing time point for each respective condition. Concentrations (pg/mL) of 13 different cytokines were measured using a LEGENDplex multiplex assay. Panels show results for: **(A)** GM-CSF, **(B)** IFN-α, **(C)** IFN-β, **(D)** IFN-γ, **(E)** IFN-λ1, **(F)** IFN-λ2, **(G)** IL-1β, **(H)** IL-6, **(I)** IL-8, **(J)** IL-10, **(K)** IL12-p70, **(L)** IP-10, and **(M)** TNF-α. Data are presented as individual biological replicates (n=3 per condition per time point) with the mean ± standard error of the mean (SEM). Statistical significance was determined by a repeated measures two-way ANOVA with Dunnett’s multiple comparison test, comparing each treatment to the time-matched VEH control. Only p-values < 0.1 are shown.

Levels of GM-CSF, IL-1β, IL-10, and IL-12p70 were either at or near indetectable (Figure 1A, G, J, K), with the absence of the immunosuppressive cytokine IL-10 proving particularly surprising, given that EBV typically upregulates the expression(47). LTA treatment induced significant upregulation in IL-6, IL-8, and IL-10 at multiple timepoints, confirming the responsiveness of HMC3 cell conditions to an inflammatory stimulus.

GP350 did not elicit substantial influences on microglial cytokine production, suggesting the glycoprotein alone is insufficient to influence microglial phenotypes and the dependency on viral particles to induce immune suppression. These results suggest that exposure to EBV particles without infection selectively suppresses the microglial IFN response, providing a putative mechanism for EBV to curate an immunosuppressive CNS TME. This selective immunosuppression may be particularly relevant in immunocompromised people, where microglia serve as the initial defense against CNS malignancies.

### UVi-EBV Regulation of Microglial Gene Expression

Next, we investigated transcriptional profiles of HMC3 cell lines activated with UVi-EBV, GP350 (Supplemental Figure 1), and LTA (Supplemental Figure 2) to investigate putative mechanisms of immune suppression. UVi-EBV exposure did not induce significant differential expression of any gene targets as detected by sequencing at 0-, 2-, 4-, or 48-hours post exposure (Figure 2A, 2B, 2C, and 2E). However, at 24-hours post UVi-EBV exposure we observed a significant downregulation in the transcriptional abundance of the pro-inflammatory immune-related genes SPP1 (ENSG00000118785), CD163 (ENSG00000177575), FCGR2A (ENSG00000143226), and PLEK (ENSG00000115956). SPP1(48, 49), CD163(50), FCGR2A(51), and PLEK(52) have each been previously indicated to possess often complex and typically pro-inflammatory roles in innate immunity, however by the 48-hour timepoint their expression levels reverted to insignificant differential expression, indicating their short term peak in expression following exposure. Interestingly, GP350 treatment downregulates SPP1, FCGR2A, and PLEK at 24-hours before a return to baseline expression at 48-hours (Supplemental Figure 1), indicating this effect observed in UVi-EBV treated samples may be a result of virion GP350. Due to the GP350 treatment, and associated expression levels of the aforementioned genes, not impacting anti-tumoral cytokine production at the 24-hour mark however (Figure 1), we infer this to not be a major mechanism of EBV-mediated immune suppression. At 72-hours post UVi-EBV exposure, we observed an upregulation in the expression of the immediate early genes (IEGs) FOS (ENSG00000170345) and EGR1 (ENSG00000120738) (Figure 2F)(53). The induction of IEGs identifies a promising molecular mechanisms for the acute changes in microglial phenotypes in response to EBV stimuli(54, 55). Our findings are complemented by previous studies which show induction of EGR1 by EBV(56, 57) and molecular mimicry between FOS and the EBV-encoded homologue BZLF1, also known as EB1(58, 59). In addition to their respective associations with EBV infection, FOS and EGR1 are well characterized proto-oncogenes with complex pro-/anti-tumoral properties(60, 61). Briefly, FOS functions as a dimeric constituent of AP-1 transcription factor complexes(62) and actively contributes to cancer promoting inflammation(63). FOS and EGR1 are primarily recognized as positive regulators of inflammation, but their impacts on inflammation are pleiotropic in nature and can also negatively regulate immunity(64–67). Next, we sought to confirm that FOS and EGR1 are co-expressed in our microglial model (Figure 2G) and are negatively correlated with supernatant concentrations of downregulated cytokines in IFNα2 (Figures 2H) and IP-10 (Figure 2I) at 72-hours. We have identified the downregulation of numerous inflammatory mediators and the upregulation of IEGs as potential mechanisms for EBV-driven suppression of microglial immunity. Collectively, our findings propose EBV-induced FOS and EGR1 signaling as potential targets mediating the suppression of microglial anti-tumor immune responses.

**Figure 2:**
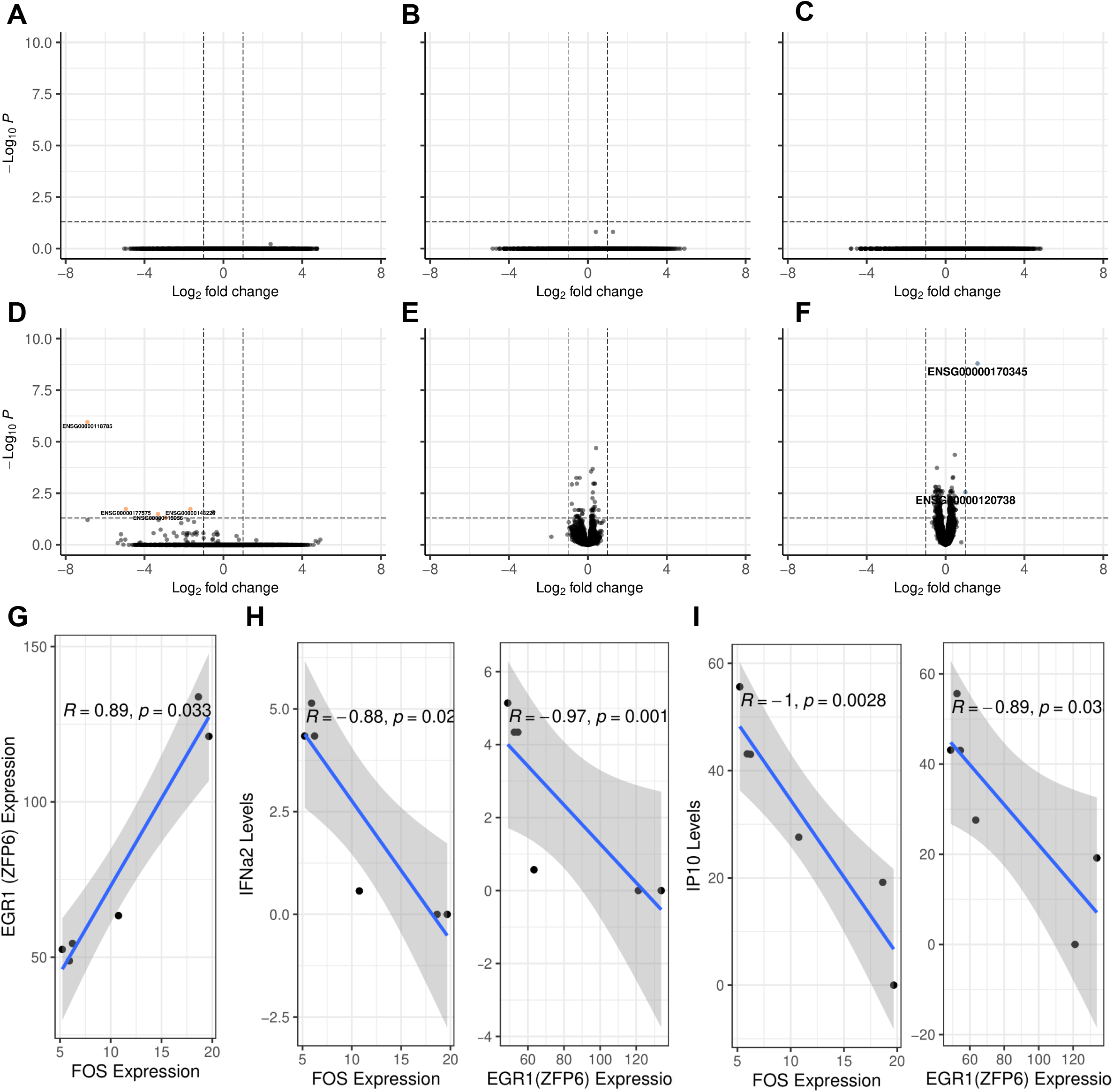
UVi-EBV Regulation of Microglial Gene Expression. Outputs of RNA sequencing from the transcriptomes of HMC3 cells treated with VEH or UV-inactivated EBV (UVi-EBV, VEH+6.4 × 10^8^ genomic copies/mL). Volcano plots exhibit differential expression patterns of human genome build 38 (hg38) annotated loci between VEH and UVi-EBV treated samples at **(A)** 0-, **(B)** 2-, **(C)** 4-, **(D)** 24-, **(E)** 48-, **(F)** and 72-hours post-exposure. Differential expression was determined with a z score using shrunken estimates of log fold change divided by standard error and a Benjamini-Hochberg correction of p values (padj). Loci were determined as differentially expressed when possessing an absolute fold change >1 and a padj <0.05. Orange dots indicate loci downregulated in UVi-EBV exposure, blue dots indicate loci upregulated in UVi-EBV exposure, and grey dots indicate unchanged loci.

## 4. Discussion

EBV is a ubiquitous gamma herpesvirus with infection rates associated with human inflammatory diseases and cancers(1, 68). Despite high infections throughout populations, EBV-associated comorbidities only develop in some individuals. This disparity between EBV infection and EBV-associated chronic disease incidences emphasize the complex nature of interactions with heterogenous populations and identifies numerous lines of inquiry for investigating host-EBV interactions. While EBV infections are strongly associated with CNS inflammation(30) and tumorigenesis(29), the mechanisms responsible for which remain poorly defined. For this purpose, we investigated the impact of EBV on human microglial immunophenotypes in a cell line based model to identify how EBV particles in the CNS may sculpt the neuroimmune environment.

Our results find that EBV exposure suppresses microglia IFN production in a time dependent manner. IFN-α2 and -β were significantly downregulated by the 72-hour time point following EBV exposure, and IFN-γ demonstrated a near-significant (padj = 0.0529) trend of downregulation. IFN-λ2 demonstrated non-detectable expression in EBV exposed cells, however the comparison were statistically insignificant due to variance in the control groups, indicating the need for further studies utilizing larger sample pools and more sensitive assays. This EBV-mediated suppression of microglia IFN production provides a putative mechanism for EBV to promote intracranial malignancies due to the key roles of IFNs in antitumor immunity(69). Mechanistically, IFNs are critical to disrupting the tumor by driving apoptosis, type 1 inflammation, and antigen presentation(70, 71). RNA sequencing analysis of these samples indicate the potential for changes in microglial IFN production levels to occur downstream or in tandem with induced expression of the IEGs FOS and EGR1. Future work on the TME of EBV+ CNS tumors may uncover rolls IEG activity in glial cells and whether their viral induction mediates pro-tumoral effects. Based on these results, we next posited that UVi-EBV exposure would yield dramatic impacts on inhibiting microglial endocytosis, an immune phenotype frequently dysregulated within the TME(72). Surprisingly, UVi-EBV had no detectable impact on microglial endocytosis (Supplemental Figure 3), suggesting compartmentalized impacts of EBV exposure on microglial biology.

This study possesses the following limitations. First, this study utilizes a cell line-based model of human microglia. This approach discounts the biological impacts of neighboring glia cells, neurons, and infiltrating immune cells on microglial phenotypes in response to EBV-derived antigen exposure. Future studies can approach these interactions by utilizing animal models, co-culturing techniques, and microglia-containing CNS organoids to determine the *in vivo* impact of EBV exposure on microglia physiology. Secondly, the utilization of a microglial cell line does not consider the heterogenous nature of microglial genotypes and phenotypes observed in nature. Interindividual heterogeneity in microglial genotypes and phenotypes possess significant impacts on host physiology(73), and therefore likely impact their responsiveness to EBV and other viral infections. In example, M1-like vs M2-like microglia may possess varying responsiveness to challenges with EBV-derived antigens. Future studies may overcome these limitations by expanding upon our findings with approaches utilizing stem cell derived microglial models(74) or primary cells derived from post-mortem tissues. Lastly, this study utilized bulk analysis of paired-end short read RNA sequencing (2×150bp). Future studies utilizing long read sequencing to identify transcript isoforms would provide more comprehensive profiling of the transcriptional activity. Furthermore, sequencing of single cells would provide additional resolution in determining the cellular biology underlying EBV-driven changes in microglial transcription and phenotypes.

In conclusion, our study demonstrates that exposure to EBV induces a distinct immunosuppressive phenotype in human microglial cell lines characterized by the inhibition of anti-tumor IFNs and the overexpression of the immediate early genes FOS and EGR1. Importantly, this EBV-driven repression of microglial IFNs provides putative mechanisms for the development of EBV+ primary CNS malignancies, particularly in the context of immunosuppressed people. These results suggest several translational opportunities for further investigation, such as IFN-replacement therapies, FOS/EGR1 pathway inhibitors, or other immunomodulatory or antiviral approaches to restore microglial antitumor surveillance in individuals at-risk or suffering from EBV-driven primary CNS malignancies. Furthermore, maturing our understanding of microglial IFN signatures with EBV infection could inform cerebrospinal fluid or circulating biomarkers for early detection of primary CNS malignancies in at-risk people. This study not only illuminates the underlying mechanics whereby EBV influences the immune landscape of the CNS but also provides a framework for reconciling its pleiotropic roles in creation of an immunosuppressive environment within PCNSLs and PILMs versus pro-inflammatory pathology seen in multiple sclerosis, thus informing future therapeutic strategies.

Given the global rise in immunocompromised populations and ubiquitous nature of EBV-infection, elucidating and developing therapeutics targeted against EBV-mediated microglial dysfunction represents a crucial, unmet need.

## Declaration of Interests

All authors have no potential conflicts of interest to disclose.

## Acknowledgments

These works are supported by US NIH grants NCI CA260691 (DFN and EC) and NIAID UM1AI164559 (DFN). The funding bodies had no influence on the planning, conduction and analysis of data performed during this study. ND and DFN conceptualized the study. DFN acquired the funding. ND and NB prepared figures for the study with help from FSR, AN, MM, and DFN. NB prepared the first draft of the manuscript with help from ND, AN, and MN. ND, NB, AN, MM, FSR, AA, EC, and DFN aided in editing manuscript drafts. AA, MM, and EC generated and provided the UVi-EBV stocks used throughout the study. ND, NB, AN, MN, and FSR performed the experimentation and analysis throughout. All authors reviewed the full text manuscript and had the final responsibility regarding decisions to submit the manuscript for publication.

## Data Availability Statement

Raw FASTQ files of bulk RNA sequencing are available at the NCBI sequence read archive (SRA) under the accession number “PRJNA1328045”.

## Supplemental Figures

**Supplemental Figure 1:** GP350 Regulation of Microglial Gene Expression

Outputs of RNA sequencing from the transcriptomes of HMC3 cells treated with VEH or GP350 (VEH+80ng/mL). Volcano plots exhibit differential expression patterns of human genome build 38 (hg38) annotated loci between VEH and UVi-EBV treated samples at **(A)** 0-, **(B)** 2-, **(C)** 4-, **(D)** 24-, **(E)** 48-, **(F)** and 72-hours post-exposure. Differential expression was determined with a z score using shrunken estimates of log fold change divided by standard error and a Benjamini-Hochberg correction of p values (padj). Loci were determined as differentially expressed when possessing an absolute fold change >1 and a padj <0.05. Orange dots indicate loci downregulated in GP350 exposure, blue dots indicate loci upregulated in GP350 exposure, and grey dots indicate unchanged loci.

**Supplemental Figure 2:** LTA Regulation of Microglial Gene Expression

Outputs of RNA sequencing from the transcriptomes of HMC3 cells treated with VEH or LTA (VEH+1μg/mL). Volcano plots exhibit differential expression patterns of human genome build 38 (hg38) annotated loci between VEH and UVi-EBV treated samples at **(A)** 0-, **(B)** 2-, **(C)** 4-, **(D)** 24-, **(E)** 48-, **(F)** and 72-hours post-exposure. Differential expression was determined with a z score using shrunken estimates of log fold change divided by standard error and a Benjamini-Hochberg correction of p values (padj). Loci were determined as differentially expressed when possessing an absolute fold change >1 and a padj <0.05. Orange dots indicate loci downregulated in LTA exposure, blue dots indicate loci upregulated in LTA exposure, and grey dots indicate unchanged loci.

**Supplemental Figure 3:** UVi-EBV Does Not Influence Microglial Endocytosis

Measures of pHrodo™ Green conjugated 10kDa Dextran beads via the FitC channel at 0-, 2-, 4-, 24-, 48-, and imaged via a 6,441μm in diameter circle (32.58mm^2^ area). The extent of microglial endocytosis was recorded in the number of endocytic vesicles (EVs) per 1mm^2^ **(A)** to quantify quantity and the sum area of captured EVs per 32.58mm^2^ **(B)** to measure quantity and size. Data are presented as individual biological replicates (n=3 per condition per time point) with the mean ± standard error of the mean (SEM). Statistical analysis of EV production was performed using a repeated measures two-way ANOVA with the Geisser-Greenhouse correction, followed by Šídák’s multiple comparison test for the generation of p-values to UVi-EBV exposure against the time-matched VEH.

## Notes

https://www.ncbi.nlm.nih.gov/bioproject/PRJNA1328045/

https://github.com/NicholasDopkins/HMC3

### Competing Interest Statement

The authors have declared no competing interest.

