## Supplementary figures and images for "EBV Triggers a Distinct Antiviral Response in HMC3 Cells"

### Supplemental Figures

#### Slide 1
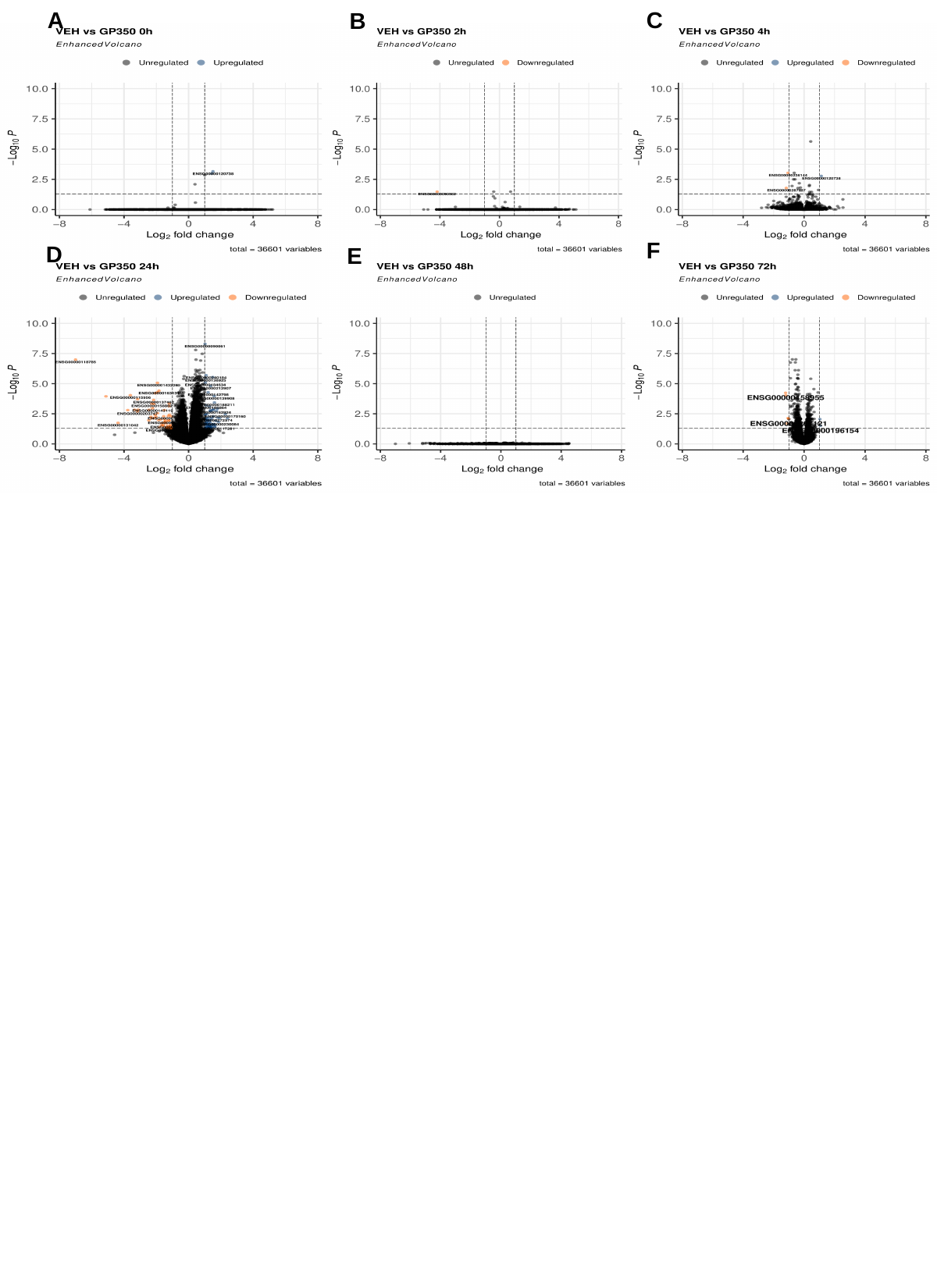

A
C
B
F
D
E

#### Slide 2
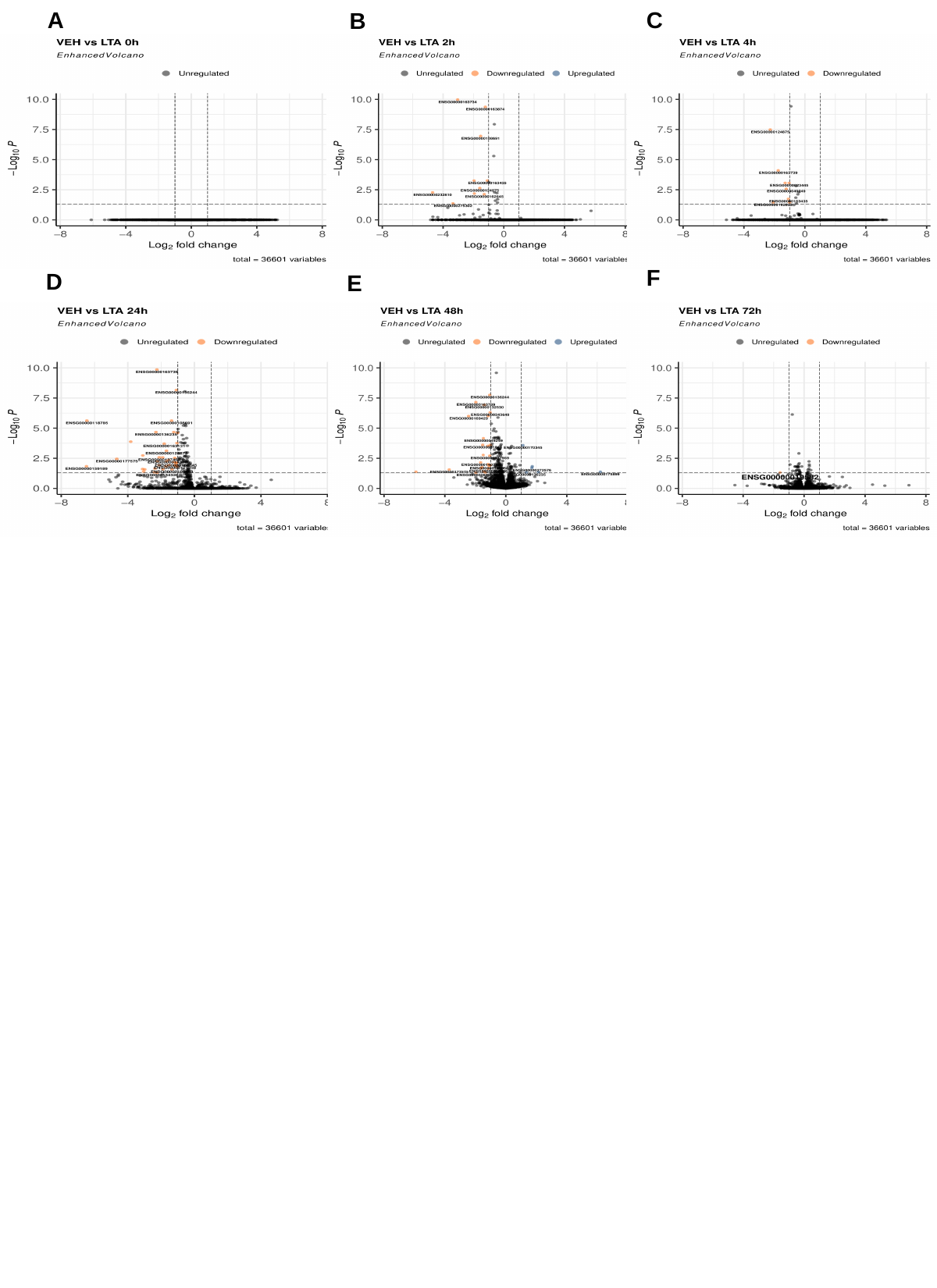

A
C
B
F
D
E

#### Slide 3
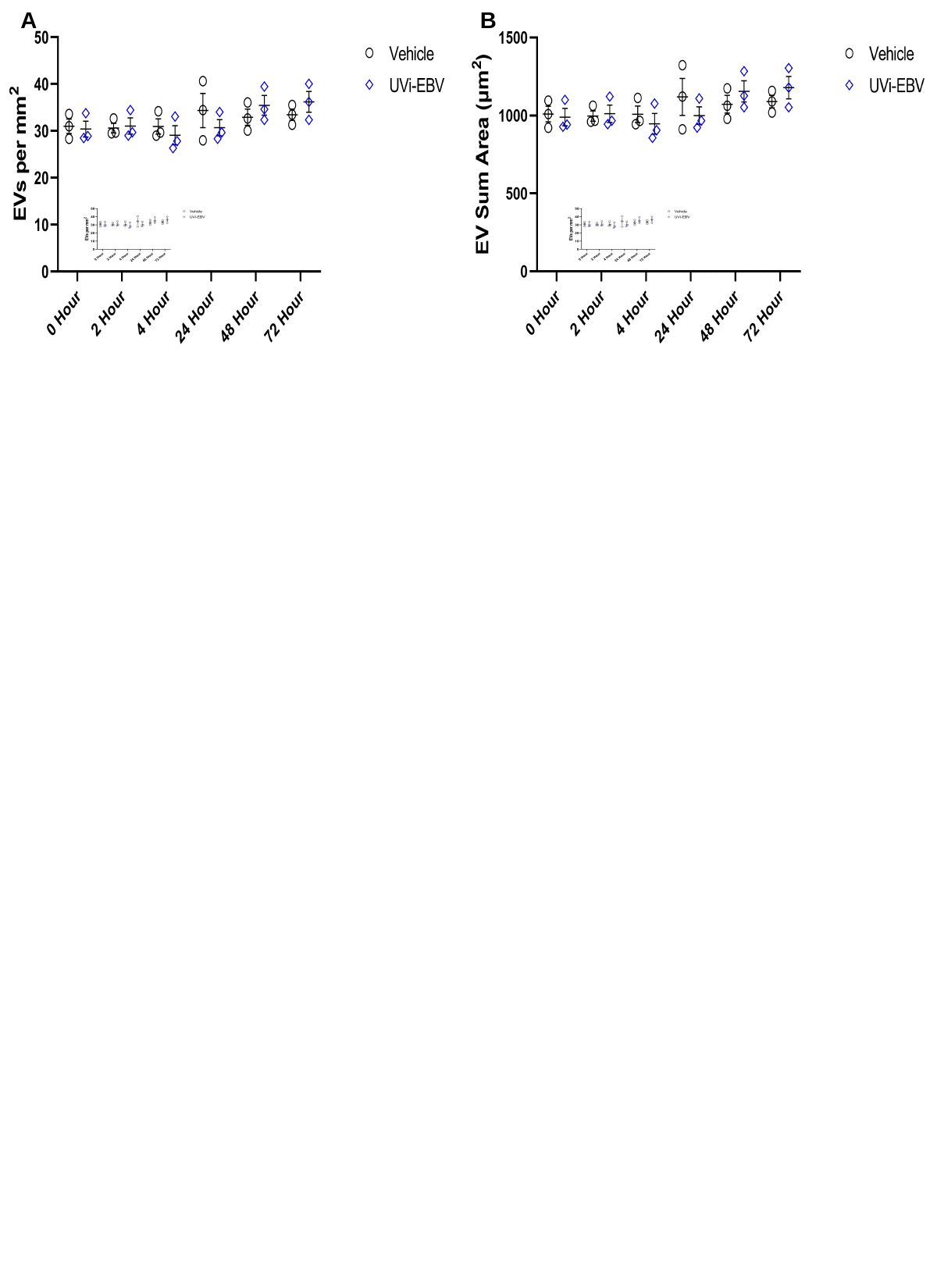

A
B
